# Fluoxetine degrades luminance perceptual thresholds while enhancing motivation and reward sensitivity

**DOI:** 10.1101/2022.11.11.516168

**Authors:** Maëva Gacoin, Suliann Ben Hamed

## Abstract

Selective serotonin reuptake inhibitors (SSRIs) increase serotonin activity in the brain. While they are mostly known for their antidepressant properties, they have been shown to improve visual functions in amblyopia and impact cognitive functions ranging from attention to motivation and sensitivity to reward. Yet, a clear understanding of the specific action of serotonin to each of bottom-up sensory and top-down cognitive control components and their interaction is still missing. To address this question, we characterize, in two adult macaques, the behavioral effects of fluoxetine, a specific SSRI, on visual perception under varying bottom-up (luminosity, distractors) and top-down (uncertainty, reward biases) constraints while they are performing three different visual tasks. We first manipulate target luminosity in a visual detection task, and we show that fluoxetine degrades luminance perceptual thresholds. We then use a target detection task in the presence of spatial distractors, and we show that under fluoxetine, monkeys display both more liberal responses as well as a degraded perceptual spatial resolution. In a last target selection task, involving free choice in the presence of reward biases, we show that monkeys display an increased sensitivity to reward outcome under fluoxetine. In addition, we report that monkeys produce, under fluoxetine, more trials and less aborts, increased pupil size, shorter blink durations, as well as task-dependent changes in reaction times. Overall, while low level vision appears to be degraded by fluoxetine, performance in the visual tasks are maintained under fluoxetine due to enhanced top-down control based on task outcome and reward maximization.

## Introduction

While selective serotonin reuptake inhibitors (SSRIs) are mostly known for their antidepressant properties (Bauer et al., 2008), serotonin concentration in the brain impacts multiple sensory and cognitive functions ranging from retinal (for review, see Masson 2019; Pootanakit and Brunken 2000) and visual functions (Lansner et al., 2019), to higher order cognitive functions such as sustained attention (Carter et al., 2005; Enge et al., 2011; Li et al., 2018; Scholes et al., 2007; Wingen et al., 2008), impulsivity (Brown et al., 2012; Meyniel et al., 2016; Worbe et al., 2014), working memory and learning (Meneses and Liy-Salmeron, 2012) and emotional and affective processing (Bar-Haim et al., 2007; Harmer, 2008; Harmer et al., 2006; Pérez-Edgar et al., 2010). Serotonin has also been proposed to play a crucial role in remodeling visual cortical circuits (Maya-Vetencourt et al. 2008; 2011; Umemori et al. 2018) as well as in cognitive flexibility (Clarke et al., 2004). In this context, the specific contribution of serotonin to low-level visual luminosity perception on the one hand and to the top-down control mechanisms of visual perception on the other hand, such as attentional selection and distractor suppression (Di Bello et al., 2022) or reward-based decision making (Homberg, 2012; Seymour et al., 2012) is still a matter of research.

Serotonin brain concentrations can be modulated by increasing the circulating levels of tryptophan, its precursor, by the action of selective serotonin agonists, or yet by the action of SSRIs. These bind selectively to serotonin transporters and inhibit their ability to reuptake serotonin into presynaptic terminals, resulting in an increase in the levels of extracellular serotonin (Wong et al., 1995; Clark et al., 1996). Thus, increased release in serotonin enhances the neurotransmitter likelihood to bind to a post-synaptic receptor. In addition, the decrease of serotonin reuptake also inhibits the negative feedback regulation, resulting in increased serotonin release in the synaptic cleft (Cerrito and Raiteri, 1979). A particular SSRI, fluoxetine (Prozac), expresses a strong binding to receptors 5-HT2c (Ni and Miledi, 1997; Pälvimäki et al., 1996) and 5-HT2a (Koch, 2002), the latter having the highest concentration in the visual system (Beliveau et al., 2017; Hansen et al., 2022), as well as in the prefrontal cortex (Puig and Gulledge, 2011). In particular, serotonin modulates the neuronal activity in the visual system, in a dose- and specie dependent-manner, such that increase or depletion of 5-HT regulates the switch between single-spike activity and rhythmic burst firing specifically in brain regions involved in visual processing, such as the retina, the visual cortex, and the thalamus (Brunken et al., 1993; McCormick and Wang, 1991; Monckton and McCormick, 2002; Moreau et al., 2013). In addition, fluoxetine decreases extracellular GABA levels (Vetencourt et al., 2008; Baroncelli et al., 2011, Beshara et al., 2016; Santana et al., 2004) thus leading to enhanced cortical excitability through a reduction of global inhibition. Relevant to the study of the effects of serotonin on top-down and bottom-up visual processes, GABAa receptor concentrations are higher in the visual cortex than in the rest of the brain and higher in the ventral part of the striate and extrastriate cortex than in its dorsal part (Kaulen et al., 2022). However, the link between GABAa receptor concentrations and cognitive readout is not straightforward. Indeed, GABA concentrations in the prefrontal cortex are negatively related to attentional blink magnitude while GABA concentrations in the posterior parietal cortex are positively correlated with attentional blink magnitude and GABA concentrations in the visual cortex do not contribute to attentional blink magnitude (Kihara et al., 2016). All this taken together indicates complex interactions between serotonin circulating levels and behavioral and cognitive markers of visual perception.

In the present work, we precisely characterize the effects of fluoxetine on visual perception under varying bottom-up (luminosity, distractors) and top-down (uncertainty, reward biases) constraints, while two monkeys perform three different visual tasks. The first task is a visual detection task in the presence of target stimuli of varying luminosity and mostly involves bottom-up visual processes. This task thus allows to characterize the effects of fluoxetine on luminosity perceptual thresholds. The second task is a visual detection task in the presence of spatial distractors and involves a combination of bottom-up visual processes and top-down target selection and reactive distractor suppression mechanisms (Di Bello et al., 2022). This task thus allows to characterize the effects of fluoxetine on perception under spatial uncertainty. The third task is a free choice task in the presence of reward biases (Chelazzi et al., 2014). This task thus allows to characterize the effects of fluoxetine on reward-based decision making. Overall, we report longer time on the task, increased pupil size and shorter blink durations under fluoxetine, increased luminance perceptual thresholds, such that higher levels of luminosity are needed to reach a 50% correct detection, more liberal decision thresholds thus producing more responses to both targets and distractors, a degraded perceptual spatial resolution under spatial uncertainty, and an enhanced sensitivity to both the positive incentive of high rewards as well as to the negative outcome of low rewards. We finally show that fluoxetine can either speed up or slow down manual reaction times, depending on the nature of the task. Overall, we show that the effects of fluoxetine on perception result from interference with both bottom-up perceptual mechanisms, namely degraded luminosity thresholds or degraded spatial resolution and top-down perceptual mechanisms, namely relaxed decision thresholds and increased sensitivity to reward outcomes. In other words, while low level vision appears to be degraded by fluoxetine, performance in the visual tasks may be maintained under fluoxetine due to enhanced top-down control based on task outcome and reward maximization.

## Results

### Under fluoxetine, monkeys work longer and produce less aborts

Based on serum concentration decay time in macaques (half-life <16h, Sawyer and Howell, 2011) and to the reported threshold for behavioral effects (Fontenot et al., 2009; Chen et al., 2012), monkeys were injected with 2.5mg/kg of Fluoxetine or an equivalent volume of saline. On each session, subjects were allowed to work for as long as they were motivated to. Monkeys were considered as less motivated and were brought back to their home cage when their compliance to the central fixation constrain in the task decreased beyond a certain threshold (85% overall fixation in a block of 280 trials for both the luminance detection task and the target detection task with distractors and 75% for the saccadic reward competition task). Note that

For all three tasks, a significant increase in the number of trials as well as a significant decrease in abort trials (i.e. trials discontinued prior to the onset of task response signal) is observed when monkeys are on fluoxetine compared to placebo sessions (Figure 1). Specifically, during the luminance detection task, both monkeys M1 and M2 performed more trials per session in fluoxetine sessions as compared to placebo sessions (M1: median number of trials per session +/-s.e.: placebo: 1257,67+/-233,98; fluoxetine: 1914,33+/-13,95; Wilcoxon non-parametric test, p=0,024 M2: placebo: 289,25+/-62,33; fluoxetine: 387+/-14,81; p=0,034) and less aborted trials (M1: median %Abort: placebo: 50,60+/-0,03; fluoxetine: 35,53+/-0,01; p=0,005; M2: placebo: 90,16+/-0,02; fluoxetine: 84,83+/-0,02; p=0,045). This was also true for the detection task with distractors (M1: median number of trials per session +/-s.e.: placebo: 970,5+/-72,85; fluoxetine: 1443,5+/-69,01; p=0,001; M2: placebo: 270,8+/-44,04; fluoxetine: 496,9+/-86,39; p=0,009; M1: median %Abort: placebo: 23,71+/-0,01; fluoxetine: 22,7+/-0,01; p=0,016; M2: placebo: 18,93+/-0,01; fluoxetine: 16,47+/-0,01; p 0,027). However, we did not observe this effect for the saccadic reward competition task (M1: median number of trials per session +/-s.e.: placebo: 936+/-109,10; fluoxetine: 1128+/-94,99; p=0,124; M2: placebo: 624+/-87,64; fluoxetine: 792+/-94,99; p=0,109; M1: median %Abort: placebo: 29,46+/-0,05; fluoxetine: 27,80+/-0,020; p=0,456; M2: placebo: 44,15+/-0,02; fluoxetine: 37,78+/-0,02; p=0,128). Overall, fluoxetine thus enhances both the motivation of the monkeys to work (more trials) as well as their compliance on the task (less aborts). Please note that, due to the nature of the tasks, performance defined as the percentage of correct trials cannot be computed.

**Figure 1:**
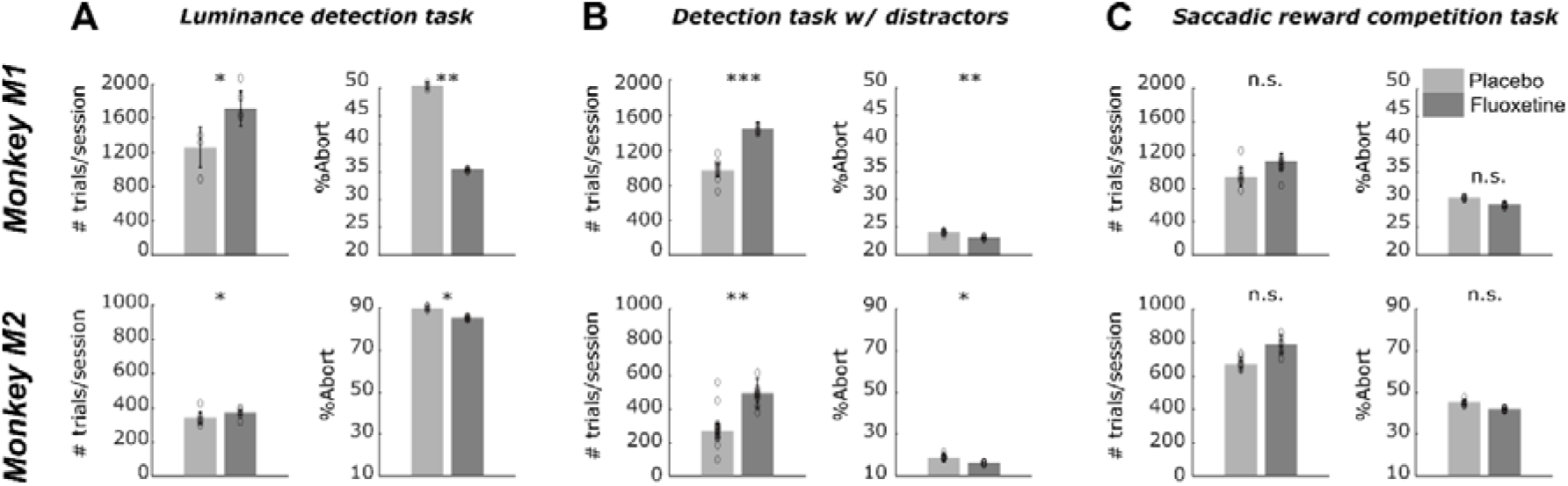
Effect of fluoxetine on on-task motivation (# of trials) and on-task compliance (% aborts), on the Luminance detection task (A), the detection task with distractors (B) and the saccadic reward competition task (C). For all plots, median +/-s.e. of the median are represented. Placebo data are represented in light gray and fluoxetine data are represented in dark gray. Statistical significance is represented as follows: ***, p<0.001; **, p<0.01; *, p<0.05; n.s., p>0.05.

### Under fluoxetine, perceptual thresholds are increased

In order to assess the effect of fluoxetine on perceptual thresholds, we had monkeys detect targets of varying luminosities ranging for very high luminance to very low, in seven steps (Figure 2). Targets could appear in one of four locations on the screen (upper left, upper right, lower left, lower right). It is to be noted that this experiment was conducted twice, at a 10 months’ interval and all observations reported below are reproduced (supplemental figure S1).

**Figure 2:**
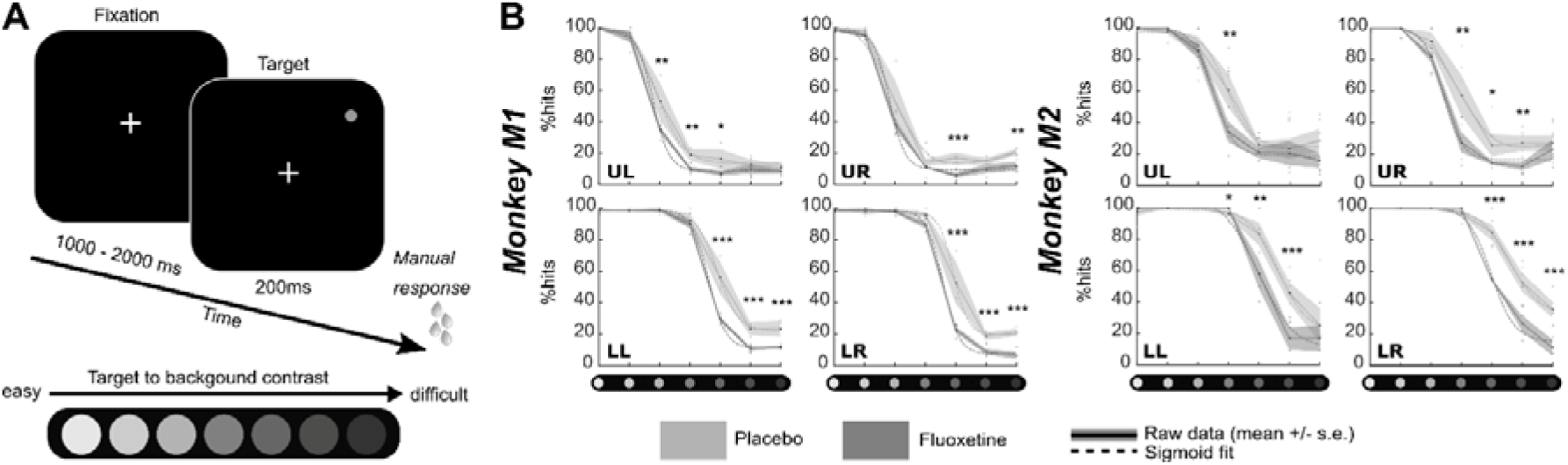
Effect of fluoxetine on perceptual thresholds in a luminance detection task. (A) Monkeys had to detect a target presented in one of four quadrants. Target luminosity ranged from high to low luminosity in 7 steps. Monkeys were rewarded for a speeded detection of target presentation. Targets were presented at four different locations, in the upper left (UL), upper right (UR), lower right (LR) and lower left (LL) quadrants, at 8° of eccentricity from the center of the screen, for 200ms. (B) For both monkeys, % of hits were computed independently for each target luminosity. Dots represent individual sessions, continuous lines represent average % hits across all sessions (+/-s.e.) and dashed lines represent sigmoid fit of the data. Placebo data are represented in light gray and fluoxetine data are represented in dark gray. Behavioral data are represented independently for each target position. Statistical significance is represented as follows: ***, p<0.001; **, p<0.01; *, p<0.05; n.s., p>0.05.

Both monkeys had a hit rate of 100% in both the placebo and fluoxetine conditions for the high luminosity targets, indicating that they were well trained and highly motivated in this task. Behavioral performance (% Hits, i.e. correct responses) were extracted as a function of target location and target luminosity for each monkey and each session and fitted with a sigmoid fit (Figure 2b, supplemental figure S1).

Two different effects of fluoxetine can be described. First, fluoxetine increases perceptual thresholds such that lower luminosity targets are less perceived under fluoxetine relative to placebo condition. This is quantified by a shift in the p^50^ (i.e. point of perceptual indecision) of sigmoid fits in the fluoxetine relative to the placebo condition towards higher luminosities (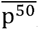, median +/-s.e. fluoxetine, M1: placebo: 4,76 ± 0,01; fluoxetine: 4,23 ± 0,04; Wilcoxon non-parametric test, p=0,026; M2: placebo: 3,90 ±0,03; fluoxetine: 3,71 ± 0,02; p= 0,002).

Second, under fluoxetine, monkeys had lower hit rates than on placebo condition, producing significantly less hits for lower luminosity targets (Figure 1B). A two-way ANOVA on target position x condition indicates a significant effect of condition and quadrant with no interaction for both monkeys (M1: target position, F(1,83)= 78,365, p<0,001; condition, F(3,249)= 77,638, p<0,001; interaction, F(3,249)= 1,639, p= 0,181; M2: target position, F(1,76)= 40,803, p<0,001; condition, F(3,228)= 41,768, p<0,001; interaction, F(3,228)= 1,702, p= 0,167). Post-hoc Wilcoxon tests indicate that this holds significant in M1 for three quadrants out of four and in M2, for one quadrant out of four (Figure 1B). This indicates that their perception for low luminosity target is strongly degraded under fluoxetine. Alternatively, this possibly indicates that monkeys become more conservative under fluoxetine, i.e. they select more carefully their responses to minimize errors. These hypotheses are evaluated in the next section, using a second behavioral task.

### Under spatial uncertainty, fluoxetine relaxes perceptual decision thresholds and degrades perceived spatial resolution

In order to further investigate the effects of fluoxetine on perceptual sensitivity and response decision thresholds described in the previous experiment, we had monkeys perform luminosity target detection task in the presence of spatial distractors (Figure 3a). In this task, targets and distractors were only distinguishable by their spatial position, and were set at a perceptual threshold of 70%, as characterized in the luminance detection task (Figure 2). Targets and distractors could either be presented in the lower left or in the lower right quadrants. As for the previous task, this experiment was conducted twice, at a 10 months’ interval and all observations reported below are reproduced.

**Figure 3:**
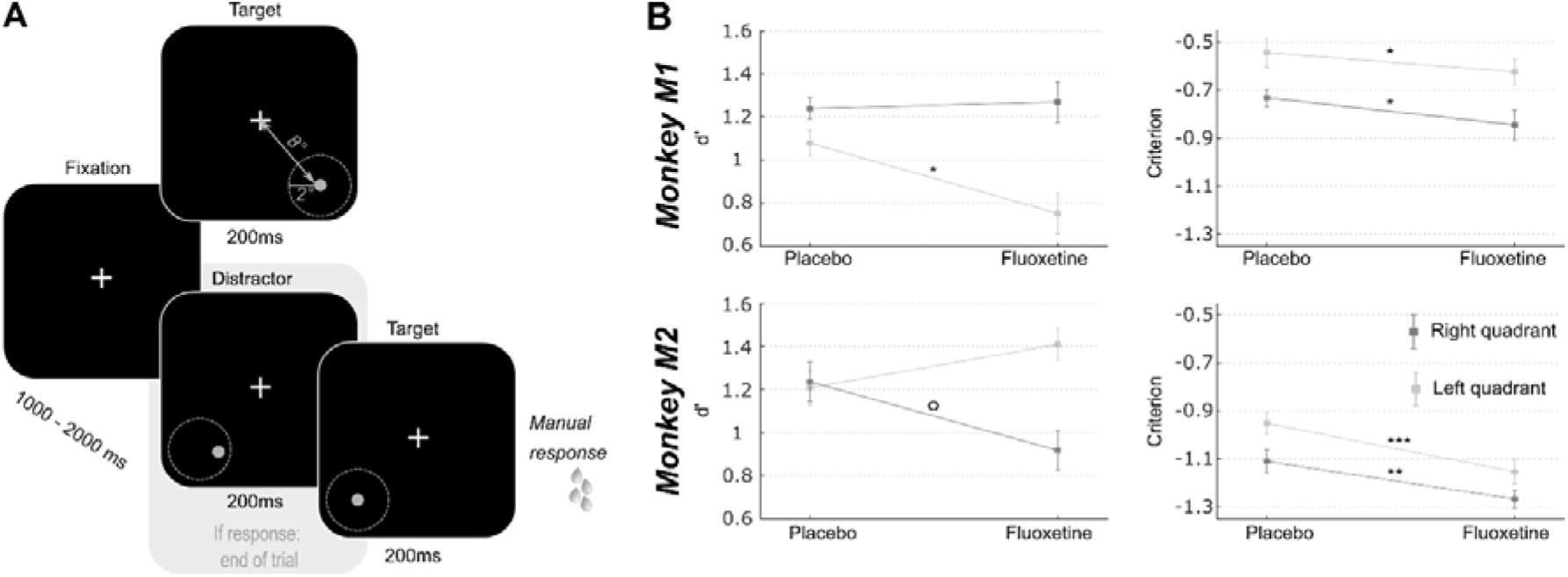
Effect of fluoxetine on spatial sensitivity d-prime and response criterion in a target detection task in the presence of spatial distractors. (A) Monkeys had to detect a target presented in one of two quadrants (lower left or lower right). Target luminosity was kept high and presented at a fixed location, at 8° of eccentricity from the center of the screen, for 200ms. Monkeys were rewarded for a speeded detection of target presentation. On 75% of the trials, targets were preceded by a distractor, undistinguishable from the target except for its spatial location. These distractors were located within a circle of 2° of eccentricity around the target. Responses to these distractors interrupted the trial and monkeys were not rewarded. (B) For both monkeys, d’ and criterion were computed independently for each target (left, light gray; right, dark gray). Median +/-s.e. of median are presented for placebo and fluoxetine conditions. Statistical significance is represented as follows: ***, p<0.001; **, p<0.01; *, p<0.05; °, p<0.07.

Based on the monkeys’ response in this task (Hits: correct target detections; Misses: no response to target presentation; False alarms: erroneous responses to distractors and Correct rejections: correct no response to distractors), we calculated the criterion (reflecting the willingness to respond that the signal is present in an ambiguous situation, independently of the subject’s sensitivity to the signal) and d-prime (reflects the actual sensitivity of the subject to the signal) over all sessions in both the fluoxetine and placebo conditions. A high criterion corresponds to a conservative behavior (i.e. less responses but mostly correct) while a low criterion corresponds to a liberal behavior (i.e. more responses but more false alarms). A high d’ indicates the signal is easily detected in the face of noise while a low d’ reflects a difficulty to detect the signal. Because reaction times differed between left and right targets, these trials were considered independently. This spatial uncertainty task was very difficult; thus, overall criteria were negative. Yet, for both monkeys and both hemifields, we observe a significant decrease in criterion after fluoxetine administration compared to the placebo condition (Figure 3b, M1: right: fluoxetine, median+/-s.e. of median, -0,84+/-0,05; placebo: -0,73+/-0,03; Wilcoxon non-parametric test, p=0,047, left: fluoxetine, -0,62+/-0,04; placebo: -0,54+/-0,06; p=0,012; M2: right: fluoxetine, -1,27+/-0,04; placebo: -1,11+/-0,05; p= 0,002, left: fluoxetine, -1,15+/-0,05; placebo: -0,95+/-0,05; p<0,001). Under fluoxetine, both subjects lowered their response decision thresholds, so that they allowed themselves more mistakes. This effect was present irrespective of whether distractors were closest to the target location, closest to the center of the visual field or further away in the periphery (supplemental figure S2a and S2b, Two-way ANOVA condition x distractor location, M1: condition, F(1,91)= 4,266, p=0,041; location, F(2,182)= 12,772, p<0,001; interaction, F(2,182)= 0,139, p=0,870; M2: condition, F(1,62)= 19,484, p<0,001; location, F(2,124)= 11,389, p<0,001; interaction, F(2,124)= 0,829, p= 0,439, this analysis cumulates both test and retest tasks to increase samples per distractor location categories). In this task, changes in d-prime (assessing sensitivity to spatial location) were inconsistent across subjects and across hemifields (M1: right: fluoxetine, median +/-s.e.: 1,27+/-0,07; placebo: 1,24+/-0,05; Wilcoxon non-parametric test, p=0,323; left: fluoxetine, 0,75+/-0,06; placebo: -0,54+/-0,06; p=0,020; M2: right: fluoxetine, 0,92+/-0,09; placebo: 1,23+/-0,09; p= 0,065; left: fluoxetine, 1,41+/-0,07; placebo: 1,21+/-0,08; p= 0,250). Because fluoxetine is expected to change excitatory/inhibitory balance in favor of inhibition through its effect on GABAergic circuitry and thus change the coding spatial resolution in the visual cortex (Robinson et al., 2003), we reasoned that changes in spatial d-primes might depend on the actual distance of the distractors to the target. For both monkeys, fluoxetine resulted in significantly decreased d-primes for close distractors but not for intermediate and far distractors (Figure S2b, intermediate distractors, M1: Wilcoxon non-parametric test, p=0,126; M2, p=0,470; close distractors, M1: p=0,041; M2, p=0,043; far distractors, M1: p = 0,155; M2: p=0,481). Post-hoc analyses however indicate that this effect was driven, in both monkeys, by a right quadrant effect. This suggests a degraded spatial resolution in the visual cortex, compatible with a related excitatory/inhibitory balance under fluoxetine.

### Fluoxetine results in increased sensitivity to reward during free choice

Decision-making in non-human primates is most often guided by reward expectation. Recent fMRI observations suggest that spatial biases induced by reward incentives are subtended by a cortical network that is functionally distinct from spatial biases induced by spatial attention (Zubair et al., 2021). Additionally, in ecological conditions, foraging often takes place in a changing environment, where the actual location of rewards change dynamically with time and the actions taken in this environment. Continuously updating expected reward locations is thus crucial. We extensively trained the two macaques included in the present study on a saccadic reward competition in a stable environment (Figure 4a). On every trial, the monkeys had to make a saccade to one of two possible targets. Each successful saccade was rewarded, but the delivered reward depended on which target was selected on this specific trial. Some targets were associated with an 80% probability of high reward and a 20% probability of low reward (High expected reward). Some targets were associated with the opposite reward contingencies: 20% probability of high reward and 80% probability of low reward (Low expected rewards). Some others yet were associated with 50% probability of high or low reward (Intermediate expected reward). Reward contingencies were fixed from one trial to the next, and were spatially organized such that each High or Low expected reward target was neighbored by Intermediate expected reward targets. Prior to our measurements, monkeys were training on the contingency map task, in a fixed reference configuration. They were then trained under a daily rotational change in this overlearned contingency map for four weeks. In order to evaluate the effect of fluoxetine on reward-based decision-making in a changing environment, we then performed acute fluoxetine (or placebo) injections while the monkeys performed the saccade reward competition task described above, to the exception that reward contingency maps varied from one day to the next in a pseudo-random manner (Figure 4b). In addition to engage monkeys in active inference of the reward contingency maps on each day, this manipulation allowed to dissociate possible effects of fluoxetine on each of reward and spatial biases. On every week, a placebo session was recorded. The next day, monkeys received an acute injection of 2,5mg/kg of fluoxetine. They then worked on the remaining weekdays on the same task, but these days were considered as washout days. Because subjects, whether human or non-human, have individual reward sensitives as well as individual spatial response biases, we independently characterize the effects of fluoxetine on reward and spatial biases as presented next.

**Figure 4:**
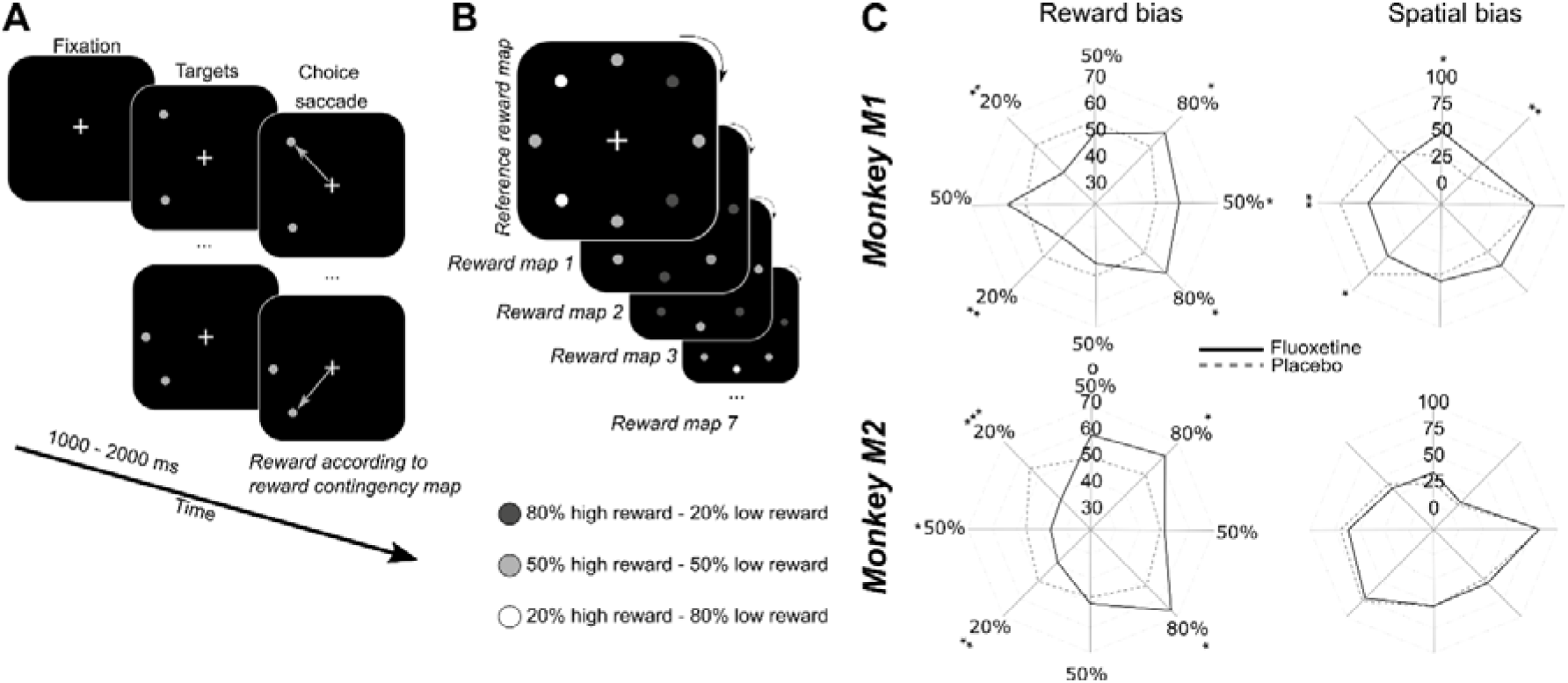
Effect of fluoxetine on saccadic choices towards targets of different reward contingencies. (A) Monkeys had to fixate a central cross on a screen 60cm away from their eyes. After an interval of 1 to 2 secs, two stimuli appeared simultaneously at two different locations out of eight. All of the 8 possible target locations were organized along a virtual circle of 8° of eccentricity from the fixation cross, equidistant one from the other. Monkeys were rewarded to make a saccadic eye movement to any of the two targets. (B) Each target was associated with two possible reward quantities, but with a different probability. High expected reward targets were associated with 80% of high reward probability and 20% of low reward probability. Low expected reward targets were associated with 20% of high reward probability and 80% of low reward probability. Intermediate expected reward targets were associated with 50% of high reward probability and 50% of low reward probability. Reward contingencies between neighbors were kept constant as follows: 80% high reward (HR) – 50% HR – 80% HR – 50% HR – 20%HR – 50% HR – 20% HR – 50% HR. However, the actual location of high and low rewarding targets changed pseudo-randomly from one day to the next. Thus, monkeys had to learn the new reward contingencies every day. We did not evaluate how much monkeys built a representation of the reference contingency map. (C) Polar plots represent the probability that monkeys choose any given target either as a function of the reward contingency map (i.e. irrespective of actual spatial position, left) or as a function of the spatial map (i.e. irrespective of actual reward contingency maps, right). Median are presented for placebo (dashed lines) and fluoxetine (continuous lines) conditions. Statistical significance is represented as follows: ***, p<0.001; **, p<0.01; *, p<0.05.

#### Effect of fluoxetine on reward biases

In order to assess the effect of fluoxetine on reward-induced biases, we computed for each individual reward contingency, a reward selectivity index (RSI) as follows. For each rewarded contingency, we estimate the median proportion of instances in which this contingency was chosen, irrespective of the reward contingency associated with the other singleton in the pair, as well as irrespective of spatial positions. Thus, RSI reflects preference for a given reward contingency irrespective of other sources of variation in the trial. Hence, a high RSI indicates that monkeys prefer this contingency relative to the others. An increase in reward selectivity index under fluoxetine indicates that the preference for this specific spatial position is enhanced. In both monkeys, we observe a significant increase in the RSI on the highly rewarded items (80% of high reward probability (M1: 80%HR_1_: fluoxetine, median +/-s.e.: 60,33+/-3,40; placebo: 52,34+/-1,72; Wilcoxon non-parametric test, p=0,028; 80%HR_2_: fluoxetine, 60,37+/-6,24; placebo: 48,22+/-1,94; p=0,044. M2: 80%HR_1_: fluoxetine, 62,62+/-4,91; placebo: 51,45+/-1,59; p=0,026; 80%HR_2_: fluoxetine, 66,44+/-5,04; placebo: 52,22+/-2,09; p=0,011) and for M1, an increase on the intermediate reward items (50% of high reward probability, M1: 50%HR_80-80_: fluoxetine, median +/-s.e.: 54,08+/-4,45; placebo: 44,90+/-0,54; p=0,0314) neighboring them (Figure 4c, left). This indicates an increase in the monkeys’ preference for these rewards under fluoxetine. We also observe, for both monkeys, a significant decrease in the RSI on both the low reward items (20% of high reward probability, M1: 20%HR_1_: fluoxetine, median +/-s.e.: 38+/-4,50; placebo: 53,85+/-2,50; p=0,005; 20%HR_2_: fluoxetine, 39,18+/-2,84; placebo: 49,97+/-2,25; p=0,006. M2: 20%HR_1_: fluoxetine, 37,15+/-4,32; placebo: 55,08+/-0,89; p<0,001; 20%HR_2_: fluoxetine, 38,88+/-3,66; placebo: 49,96+/-1,74; p=0,009) and for M2 on the intermediate reward items (50% of high reward probability, M2: 50%HR_20-20_: fluoxetine, median +/-s.e.: 36,24+/-5,16; placebo: 46,18+/-1,88; p=0,048) neighboring them (Figure 4c). This indicates a decrease in the monkeys’ preference for these rewards under fluoxetine. Thus, overall, this demonstrates that fluoxetine significantly alters reward–based decision making such that subjects are more sensitive to the positive incentive of high reward probabilities as well as to the negative outcome of low reward probabilities.

#### Effect of fluoxetine on spatial biases

In order to assess the effect of fluoxetine on intrinsic spatial biases, we computed for each individual target position, a spatial selectivity index (SSI) as follows. For each spatial location, we estimated the median proportion of instances in which this position was chosen, irrespective of the spatial position of the other singleton in the pair, as well as irrespective of reward contingencies. Hence, a high SSI indicates that monkeys prefer this contingency relative to the others. An increase in spatial selectivity index under fluoxetine indicates that the preference for this specific reward contingency is enhanced. Under fluoxetine, Monkey M1 shows a decreased SSI specifically for the left targets relative to the placebo and an increased SSI in upper positions (Figure 4c, right). Because spatial positions on the left hemifield were associated with a high SSI in the placebo condition relative to the right targets, this indicates that the monkeys often preferred targets on this side on the placebo condition and that this spatial bias decreased under fluoxetine (M1: Middle-left: fluoxetine, median +/-s.e.: 47,39+/-6,13; placebo: 77,36+/-6,78; Wilcoxon non-parametric test, p=0,003; Low-left: fluoxetine, 50,34+/-7,99; placebo: 76,58+/-8,21; p=0,020). Likewise, Monkey M1 had a low SSI toward upper positions in placebo position relative to lower positions targets and this spatial bias decreased under fluoxetine (M1: Up: fluoxetine, median +/-s.e.: 49,29+/-7,65; placebo: 22,99+/-8,51; p=0,020; Up-right: fluoxetine, 33,07+/-5,38; placebo: 13,29+/-3,37; p=0,004). Thus, in this monkey, fluoxetine resulted in a reduction in overall spatial biases. Monkey M2 show no significant difference in SSIs between the placebo and the fluoxetine condition (Figure 4c, right).

### Pupil size is enlarged and blink duration decreased under fluoxetine

Changes in perceptual thresholds as described in the first luminance detection task can be accounted for by local changes in excitatory/inhibitory balance in the visual cortex. However, this can also be accounted for by changes in oculomotor functions such as pupil size changes and blink duration (LeDoux et al., 1998). We thus quantified these two parameters independently in the fluoxetine and placebo conditions.

Overall, pupil size significantly increases under fluoxetine relative to placebo, both at rest and in the task (Figure 5A, two-way ANOVA, condition x epoch: M1: main condition effect, F(1,2999)= 1910, p<0,001; main epoch effect: F(1,2999)= 64926, p<0,001; interaction: F(1,83)= 6027, p<0,001; M2: main condition effect, F(1,2999)= 58539, p<0,001; main epoch effect: F(1,2999)= 57, p<0,001; interaction: F(1,2999)= 55, p<0,001). This observation is in agreement with what has already been described in the literature (Cazettes et al., 2021; McGuirk and Silverstone, 1990). However, enlarged pupil size is associated with enhanced visual acuity (Leibowitz, 1952). Thus, this observation on pupil size is at odds with our observation of degraded perceptual thresholds in the luminance detection task.

**Figure 5:**
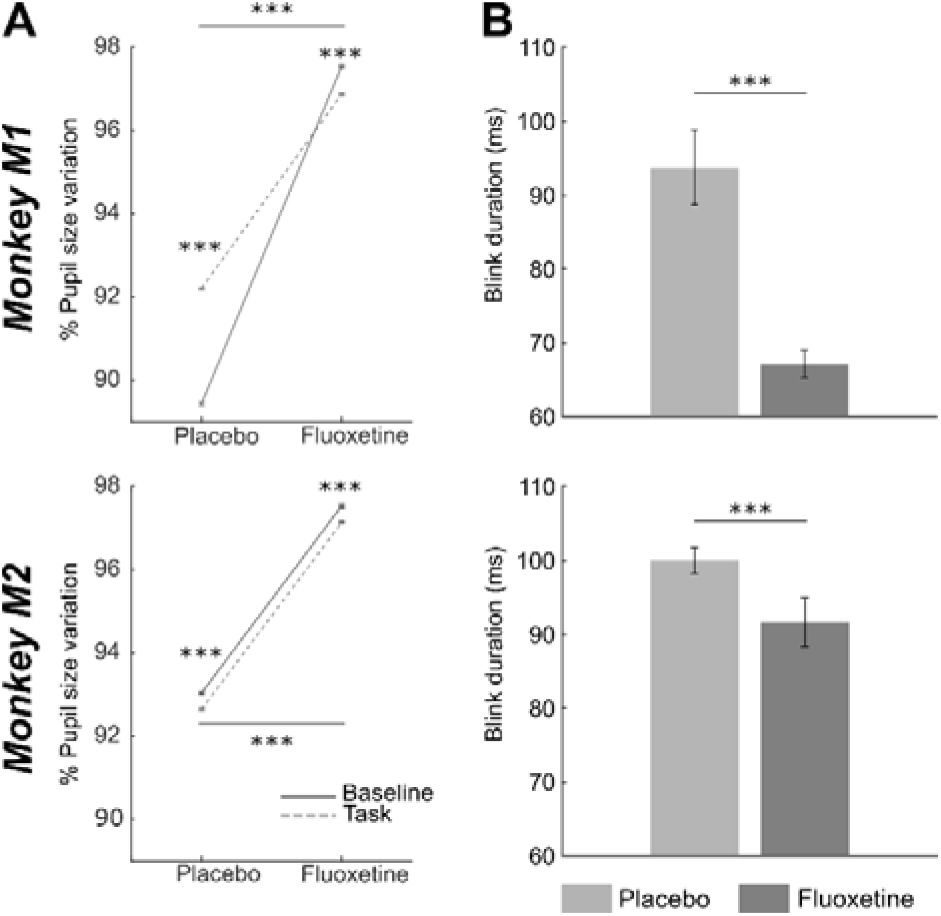
Effect of fluoxetine on pupil size and eye blink duration. (A) Median +/-s.e. of %pupil size change relative to the entire task, for monkeys M1 (top) and M2 (bottom), during rest (baseline, continuous line) and after target presentation (task, dashed lines), in the placebo and fluoxetine conditions. (B) Median +/-s.e. of blink duration, for monkeys M1 (top) and M2 (bottom), in the placebo and fluoxetine conditions. Statistical significance is represented as follows: ***, p<0.001; **, p<0.01; *, p<0.05.

We also measured eye blink statistics as a proxy of attentional engagement. Blink rate in the task was not affected by fluoxetine (M1: Median+/-s.e., fluoxetine= 0,18+/-0,001 blinks/sec; placebo= 0,19+/-0,02 blink/sec; Wilcoxon test, p= 0,152; M2: fluoxetine= 0,13+/-0,01 blink/sec; placebo= 0,13+/-0,01 blink/sec; Wilcoxon test, p= 0,290). However, blink duration was significantly shorter under fluoxetine relative to placebo (Figure 5B, Wilcoxon test, M1: p<0,001; M2: p<0,001). Because epochs of blinking have been shown to interfere with cognition (Irwin, 2014), shorter eye blink under fluoxetine might be associated with stronger involvement in the task.

### Paradoxical effects of fluoxetine on manual reaction times

Reaction times (RT) correspond to a complex behavioral variable that is subject to modulations by multiple cognitive functions ranging from spatial attention (Wardak et al., 2011, 2012a), to temporal expectation and anticipation (Cravo et al., 2013; Wardak et al., 2012b), decision making (Fujimoto et al., 2021; Hanks et al., 2006; Noorani and Carpenter, 2016), perception (Song et al., 2008), reinforcement learning (Viejo et al., 2018), arousal (Davranche et al., 2006; Eason et al., 1969; Fujimoto et al., 2021), reward (Epstein et al., 2011; Firestone and Douglas, 1975; Procyk et al., 2000; Simen et al., 2009), to name a few. In the following, we characterize the effect of fluoxetine on RT distributions. For the sake of clarity, in the following, we focus on manual reaction times in the two first tasks, as saccadic reaction time from the saccadic reward choice task are confounded by possible spatial and reward biases. We used the LATER model in order to classify reaction times in anticipatory reaction times and controlled reaction times (supplemental figure S3).

For the luminance detection task, because reaction times vary as a function of target luminance, we focused on the trials with two highest target luminance. On these trials, both monkeys had a 100% hit rate. We pooled the trials corresponding to the two easiest targets on all four positions. We report, in both monkeys, a significant decrease in controlled reaction time under fluoxetine relative to placebo (Figure 6, M1: median RT +/-s.e., placebo = 526,4ms +/-4,38; fluoxetine = 506,4ms +/-3,21; p < 0,001. M2 median RT +/-s.e., placebo = 497,1ms +/-6,19; fluoxetine = 485,9 +/-5,49; p<0,001). Fluoxetine did not have the same impact on the rate of anticipatory responses in each monkey. M1 had fewer anticipations in the fluoxetine condition (2,89%) relative to the placebo condition (7,83%, p<0,001) and faster responses in the fluoxetine condition (median RT +/-s.e., 263,8ms +/-2,99) relative to the placebo condition (329,3ms +/-4,30; p<0,001). In M2, anticipation rate was not significantly different in the fluoxetine condition (7,08%) relative to the placebo condition (6.06%, p=0,344) and these anticipatory responses were slower in the fluoxetine condition (median RT +/-s.e., 286,7ms +/-6,16) relative to the placebo condition (234,4ms +/-4,67; p=0,029).

**Figure 6:**
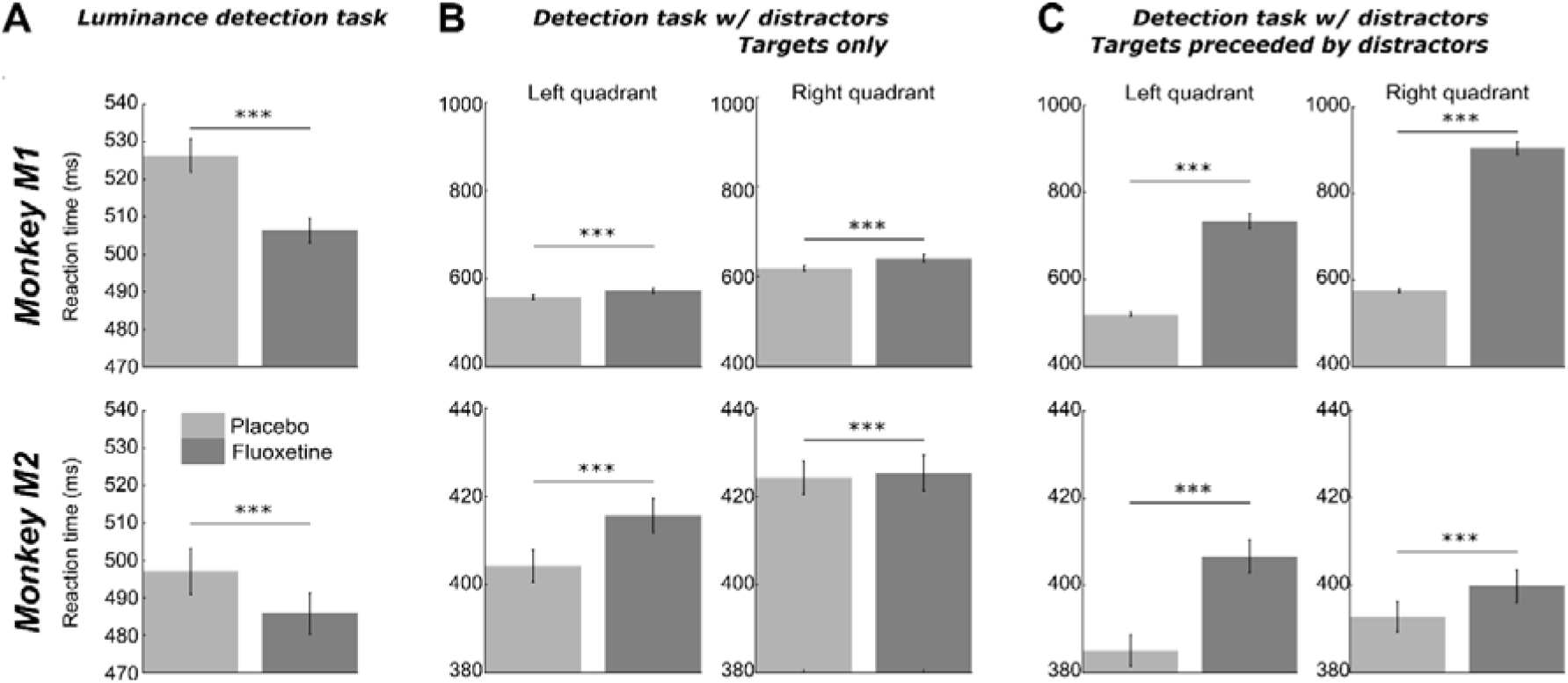
Effect of fluoxetine on controlled reaction times. Median +/-s.e. of reaction times, for monkeys M1 (top) and M2 (bottom), in the placebo and fluoxetine conditions in the luminance task (A, all positions collapsed, high luminance targets only), in the detection tasks with distractors, on target only trials (B), and in trials with distractors (C). Statistical significance is represented as follows: ***, p<0.001; **, p<0.01; *, p<0.05.

For the target detection task in the presence of distractors, we report the opposite observations. Indeed, RT increased under fluoxetine in target only trials (M1: median RT +/-s.e., placebo_RIGHT_ = 619,3ms +/-5,77; placebo_LEFT_ = 557,9ms +/-5,24; fluoxetine_RIGHT_ = 641,1ms +/-8,29; fluoxetine_LEFT_ = 572,5ms +/-6,90; pPLACEBO-FLUOX <0,001. M2 median RT +/-s.e., placebo_RIGHT_ = 424,4ms +/-3,80 ; placebo_LEFT_ = 404,3ms +/-3,62; fluoxetine_RIGHT_ = 425,4ms +/-4,03; fluoxetine_LEFT_ = 415,7 ms +/-3,91; pPLACEBO-FLUOX < 0,001). On these trials, we also report less percentage of anticipatory responses under fluoxetine relative to placebo (M1, left quadrant, placebo = 3,62%, fluoxetine = 1,25%, p <0,001; right quadrant, placebo = 8,86%, fluoxetine = 1,25%, p= 0,008; M2, left quadrant, placebo = 1,79%, fluoxetine = 0,55%, p<0,001; right quadrant, placebo= 0,92%, fluoxetine=0,59%, p<0,001).

On trials with a distractor preceding target presentation, RT also increased under fluoxetine in target only trials (M1: median RT +/-s.e., placebo_RIGHT_ = 575,5ms +/-5,50; placebo_LEFT_ = 520,15ms +/-4,78; fluoxetine_RIGHT_ = 906ms +/-14,46; fluoxetine_LEFT_ = 733,7ms +/-16,23; pPLACEBO-FLUOX < 0,001. M2 median RT +/-s.e., placebo_RIGHT_ = 392,8ms +/-3,49 ; placebo_LEFT_ = 385,1ms +/-3,52; fluoxetine_RIGHT_ = 399,8ms +/-3,70; fluoxetine_LEFT_ = 406,7 ms +/-3,85; pPLACEBO-FLUOX < 0,001). In addition, we observe a marked increase of the overall percentage of anticipatory responses (M1, left quadrant, placebo = 25,74%, fluoxetine = 91,75%, p<0,001; right quadrant, placebo = 47,49%, fluoxetine = 97,74%, p<0,001. M2, left quadrant, placebo = 9,71%, fluoxetine = 10,06%, p<0,001; right quadrant, placebo= 3,37%, fluoxetine=25,48%, p<0,001), and significantly more when distractor is preceding the target compared to target only trials under fluoxetine than in the placebo condition (Two-way ANOVA condition distractor presence, M1: left quadrant, condition, F(1,8)= 3586,177, p<0,001; location, F(1,8)= 1123,003, p<0,001; interaction, F(1,8)= 857,459, p<0,001; right quadrant, condition, F(1,8)= 3586,177, p<0,001; location, F(1,8)= 1123,003, p<0,001; interaction, F(1,8)= 857,459, p<0,001 ;M2: left quadrant, condition, F(1,15)= 3586,177, p<0,001; location, F(1,15)=1123,003, p<0,001; interaction, F(1,15)= 857,459, p<0,001; right quadrant, condition, F(1,15)= 3586,177, p<0,001; location, F(1,15)= 1123,003, p<0,001; interaction, F(1,15)= 857,459, p<0,001). Overall, on this task, we thus report a paradoxical effect of fluoxetine, associated with more anticipatory RTs on distractor trials, indicating a stronger release of proactive inhibitory mechanisms (Wardak et al., 2012a, 2012b; Criaud et al., 2012), while at the same time we report longer controlled RT on these same trials, indicating stronger cognitive control.

## Discussion

In the present work, we precisely characterize the effects of fluoxetine on behavioral and physiological metrics while monkeys are performing three different visual tasks. We report a set of specific effects of fluoxetine as well as several non-specific effects of fluoxetine, including longer time on the task and shorter blink durations. Luminance perceptual thresholds are increased, such that higher levels of luminosity are needed to reach a 50% correct detection. Under sensory uncertainty, decision thresholds are released and perceptual spatial resolution is degraded. We additionally show that fluoxetine increases sensitivity to reward outcome. Last, we show that fluoxetine can either speed up or slow down manual reaction times, depending on the nature of the task. In the following, we discuss these observations in the light of the current knowledge on fluoxetine.

### Fluoxetine interferes with retinal functions

We here show that fluoxetine results in increased visual perceptual thresholds, higher levels of luminosity being required to achieve similar detection thresholds as in placebo. This can be accounted for by the reported role of serotonin (5-hydroxytryptamine or 5-HT) in the physiology and development of the retinal of vertebrates (for review, see Masson, 2019; Pootanakit and Brunken, 2000). 5-HT is synthesized as a precursor for melanopsin in both photoreceptors and amacrine cells (Millar et al., 1988; Pourcho, 1996; Vaney, 1986) and its uptake occurs in bipolar and retinal ganglion cells (RGC). In particular, amacrine cells are major transporters of rod signals to RGC, playing a role in increasing their slow potential information (Nelson, 1982; Smith and Vardi, 1995). Accordingly, in fish, fluoxetine enhances serotonin accumulation in bipolar cells (Schuette and Chappell, 1998). A similar 5-HT uptake is described in cat RGC following the injection of a serotoninergic neurotoxin (Wassle et al., 1987). This results in the suppression of spontaneous spike firing in RGC, thus decreasing melanopsin driven response (Hughes et al., 2016). Overall, the coupled retinal effect of the SSRI on decreasing melanopsin and the rods transporter activity of amacrine cells is thus responsible for luminance perception depletion, and possibly accounts for our experimental observation of a p^50^ indecision threshold shifted toward higher luminosities.

### Fluoxetine interferes with pupil and blink physiology

SSRIs have been shown to result in increased pupil dilation (Hughes et al., 2016; Schmitt et al., 2002). In particular, patients treated with fluoxetine have bigger pupillary diameters and slower pupillary contraction (Rodriguez et al., 2020). It is unclear whether these effects are also associated with low level changes in visual accommodation (Rodriguez et al., 2020). Enlarged pupil diameter has been associated with higher thresholds at detecting the frequency at which a flickering light is perceived as a steady light source (Schmitt et al., 2002) as well as with enhanced letter identification report at very short presentation timings (Lansner et al., 2019). This suggests larger pupil size impacts perception and possibly also accounts for our experimental observation of a p^50^ indecision threshold shifted toward higher luminosities.

To our knowledge, there are no reports that fluoxetine impacts blink duration. Capitão et al., (2015) show that fluoxetine modulates emotional processing, suppressing for example the motion-potentiated startle effect. Here, we show a significant decrease in blink duration, in the absence of change in blink frequency. In the context of our task, these blinks are considered as spontaneous rather than reflex blinks in response to external events. Spontaneous blinks have been shown to correlate with the activation of a network involving somatosensory primary and secondary areas, as well as parietal, cingulate, insular, as well as striate and extrastriate visual areas (Guipponi et al., 2015). Shorter blinks possibly correlate with weaker activations in this network. This remains to be explored as well as the possible link between blink duration and perception.

### Fluoxetine enhances motivation on task

Reward is a key motivational factor and reward processing is known to regulate cognitive functions such as attention, memory, decision making and learning (Arnsten and Rubia, 2012; for review, see Hélie et al., 2017). Fluoxetine is proposed to mediate these cognitive functions through the reward valuation pathways. Indeed, Inhibition of central serotonin reuptake decreases probabilistic learning (Chamberlain et al., 2006) and SSRI enhances reward processing in healthy adults (Macoveanu, 2014; McCabe et al., 2010; Scholl et al., 2017). The serotoninergic cells of the dorsal raphe nucleus project to both the bed nucleus of the stria terminalis, an anxiety-related structure, the ventral tegmental area, a reward-related structure, and are shown to respond to emotional salience (Paquelet et al., 2022). The activity of this neuronal population is additionally shown to correlate with learning rate, both in a context of expected and unexpected uncertainty (Grossman et al., 2022). Fluoxetine has also been associated with a reduction of effort cost, or to an increased valuation of reward. Indeed, a study in healthy humans shows that subjects who received a SSRI put more effort to get a reward (Meyniel et al., 2016). Last, fluoxetine has been shown to decrease both hunger and thirst (McGuirk and Silverstone, 1990). In spite of this convergent evidence for a role of serotoninergic pathway on motivation through reward processing and valuation, caution is required as it has been shown that changes in sensitivity to the reward under SSRI is highly dose-dependent (Bari et al., 2010).

We report that monkeys make more trials and produce less abort trials under fluoxetine relative to the placebo condition. These observations are reproduced in three different behavioral tasks and both monkeys. This could be accounted for by an enhanced motivation to work. This enhanced motivation could be due to increased hunger or thirst. This however seems unlikely, as in humans, fluoxetine has been shown to decrease hunger and thirst rather than increase them (McGuirk and Silverstone, 1990). A general effect of fluoxetine on effort and reward valuation is more plausible (Meyniel et al., 2016). Accordingly, we observe that both monkeys expressed, in addition to an increase in the number of task trials, a higher willingness to initiate working sessions, at all stages of experimental preparation and execution (higher willingness to come out of the cage, and go in monkey chair, faster eye calibration, no signs of restlessness at the end of the working session that would indicate that the monkey wants to go back to its home cage).

In the free choice task, we manipulated reward contingency and we measured the monkeys’ sensitivity to reward. Fluoxetine significantly altered reward based-decision making such that subjects were more sensitive to the positive incentive of high reward probability as well as to the negative outcome of low reward probabilities. In other words, monkeys’ decision-making was more impacted by expected reward under fluoxetine. Thus, not only did monkeys put more effort to get a reward under fluoxetine (Meyniel et al., 2016), but they also better used reward information in order to guide their behavior.

### Fluoxetine interferes with attention

Shorter blinks (Hsieh and Tai, 2013) as well as enlarged pupil (Aston-Jones and Cohen, 2005; Wang et al., 2018) have been associated with higher arousal. Improved arousal could thus be at the origin of the enhanced commitment to the task. Beyond this non-specific arousal effect, enhanced performance in task could also be due to enhanced attention, taking place independently from motivational factors. Indeed, it has been shown that 5-HTP (the immediate serotonin precursor) uptake increases attention in low baseline attention individuals (Weinberg-Wolf et al., 2018). Likewise, fluoxetine is shown to selectively modulate, prefrontal synaptic growth during macaque brain development (Golub et al., 2017). It is also shown to activate other cortical structures involved in sustained attention, such as the thalamus and caudate nucleus in healthy subjects but no behavioral effect on sustained attention has been reported (Wingen et al., 2008) and the dorsolateral prefrontal cortex in attention deficit hyperactivity disorder and autism spectrum disorder patients (Chantiluke et al., 2015; Hollander et al., 2005; Quintana et al., 2007; Riggs et al., 2007; Strawn et al., 2015). Last, frontoparietal inhibitory mechanisms have been shown to be closely linked with individual differences in attentional processing such that GABA concentrations in the prefrontal cortex are negatively related to attentional blink magnitude while GABA concentrations in the posterior parietal cortex are positively correlated with attentional blink magnitude (Kihara et al., 2016). In other words, the functional roles of the GABAergic system in selective attention differ between the prefrontal and the parietal cortex. In contrast, GABA concentrations in the visual cortex do not contribute to neither attentional blink magnitude nor to first-target accuracy during a rapid serial visual presentation (Kihara et al., 2016). The GABA-mediated effects of fluoxetine are complex. Indeed, fluoxetine increases the cerebrospinal fluid GABA levels (Gören et al., 2007), indirectly affecting the GABA levels in the brain (Beshara et al., 2016; Santana et al., 2004). This results in a change in the excitatory/inhibitory balance in the brain to the benefit of a stronger inhibition (Yin et al., 2021). However, all this taken together suggests a specific impact of fluoxetine on the fronto-parietal attentional network (Ibos et al., 2013). A direct role of fluoxetine on the correlated activation of the fronto-parietal attentional network is observed in the same macaques as those included in the present study, during the performance of a perceptual task during an fMRI protocol (Gacoin, PhD dissertation). This thus confirms the impact of fluoxetine on the cortical substrates of the attentional function.

Attentional control on perception involves both changes in perceptual sensitivity and changes and the decision response threshold. The neuronal activity in the prefrontal cortex has been associated to both (Luo and Maunsell, 2018), while the neuronal activity of extrastriate cortex has mostly been associated with changes in sensitivity (Martinez-Trujillo and Gulli, 2018). Here, in a spatial decision task involving a spatial uncertainty, we report both a change in response criterion, monkeys becoming more liberal, as well as a loss of spatial resolution in visual processing. In line with the observations by Kihara et al. (2016), we predict that GABA concentrations in the prefrontal cortex are negatively related to response criterion magnitude, thus accounting for our experimental observations under fluoxetine. This predicts a decrease in functional connectivity between the prefrontal and parietal cortex under fluoxetine.

### Fluoxetine and reaction times

Reaction times (RT) are modulated by multiple cognitive functions ranging from spatial attention (Wardak et al., 2012a, 2011), to temporal expectation and anticipation (Cravo et al., 2013; Wardak et al., 2012b), decision making (Fujimoto et al., 2021; Hanks et al., 2006; Noorani and Carpenter, 2016), perception (Song et al., 2008), reinforcement learning (Viejo et al., 2018), arousal (Davranche et al., 2006; Eason et al., 1969; Fujimoto et al., 2021) and reward processing (Epstein et al., 2011; Firestone and Douglas, 1975; Procyk et al., 2000; Simen et al., 2009). RTs have recently been found to be prolonged in the context of decision-making under SSRIs (Khalighinejad et al., 2022). We here reproduce this observation (prolonged RTs under fluoxetine) in a detection task under spatial uncertainty as well as in a task involving free choice based on reward incentives. However, we report the opposite trend (i.e. speeded up RTs under fluoxetine) in a luminance detection task. We propose to interpret this paradoxical effect in the context of stochastic resonance. A recent study by (Groen et al., 2018) shows that adding noise to the visual cortex using transcranial random noise stimulation enhanced decision-making when stimuli were just below perceptual threshold, but not when they were well below or above threshold. Stretching this observation, we would like to propose that the observed paradoxical effects of fluoxetine on RT depend on the specific noise functions associated with each task, noise being defined as both neuronal noise possibly effected by fluoxetine due to changes in the excitatory/inhibitory balance in the brain (Yin et al., 2021), as well as task related noise or uncertainty, be it spatial uncertainty or reward-related uncertainty.

### Fluoxetine and cognitive flexibility

Cognitive flexibility corresponds to the ability animals have to dynamically change their behavioral strategy as a function of the external contingencies as well as their internal drives. Prefrontal serotonin levels have been associated with enhanced behavioral flexibility (Clarke et al., 2004). Here, we show that under fluoxetine, in a free choice task in which reward contingencies change daily and thus have to be relearned, fluoxetine resulted in a stronger implementation of reward outcome in the choice strategy of the monkeys. This thus indicates that monkeys were more flexible in adapting their response choice from one day to the next and from one trial to the next, confirming an enhanced level of cognitive flexibility.

### GABA-, Dopamine- and noradrenaline-mediated effects of fluoxetine on perception

Perception relies on both lower level visual mechanisms and higher order attentional mechanisms. As discussed in a previous section, frontoparietal inhibitory mechanisms are closely linked with individual differences in attentional processing (Kihara et al. 2016), such that the functional roles of the GABAergic system in selective attention differs between the prefrontal and the parietal cortex. In contrast, GABA concentrations in the visual cortex do not contribute to neither attentional blink magnitude nor to first-target accuracy during a rapid serial visual presentation (Kihara et al. 2016). It is unclear whether visual cortex GABA concentrations affect perceptual luminosity thresholds independently of top-down attentional processes. GABAa receptor distribution is not homogenous throughout the brain (Kaulen et al., 2022). In particular, GABAa receptor concentrations are higher in the ventral part of the striate and extrastriate cortex than in its dorsal part (Kaulen et al., 2022). Due to this GABAa inhomogeneity, predicting the cognitive effects of systemic fluoxetine neuromodulation is complex. For example, this bias in GABAa receptor concentrations between dorsal and ventral occipital cortex possibly contributes the well documented functional asymmetry between the upper and lower visual fields (Carlsen et al., 2007, 2007; Chen et al., 2005; Khan and Lawrence, 2005; Liu et al., 2006; Maehara et al., 2004; Qu et al., 2006; Rizzolatti et al., 1987) and higher spatial resolution in the lower visual field (Levine and McAnany, 2005; Sample et al., 1997). This asymmetry has been proposed to have an evolutionary origin (Previc, 1990) due to the fact that hand and tool manipulation mostly occurs in the lower visual field (Levine and McAnany, 2005). In our own data, the luminance detection task, we observe that under fluoxetine, monkeys produce less responses to very low luminosities, specifically in the lower visual field (which is coded by the dorsal part of the occipital cortex). We propose that in the upper visual field, due to the higher GABAa receptor concentrations in the ventral part of the striate and extrastriate cortex, noise cancellation is efficient including in the absence of fluoxetine. Due to the lower GABAa receptor concentrations in the ventral part of the striate and extrastriate cortex, noise cancellation in this part of the visual field is enhanced under fluoxetine.

This highlights the complexity of predicting the specific effects of fluoxetine on cognition. To bring further complexity to this question, at high doses, fluoxetine has been shown to enhance synaptic dopamine and noradrenaline in the rat prefrontal cortex and in the hypothalamus (Bymaster et al., 2002; Itti and Koch, 2001; Pälvimäki et al., 1996; Perry and Fuller, 1997; Pinna et al., 2009), influencing the mesocortical dopaminergic pathway (Gobert et al., 2000). Overall, we show that fluoxetine interferes with bottom-up perceptual mechanisms, namely degraded luminosity thresholds and degraded spatial resolution as well as on top-down perceptual mechanisms, namely relaxed decision thresholds and increased sensitivity to reward outcomes. In other words, fluoxetine degrades low level visual functions while at the same time maintaining performance in the visual tasks due to enhanced top-down control based on task outcome and reward maximization.

## Material and methods

### Animals and ethical approval

Two healthy adult male rhesus macaques (*macaca mulatta*) took part in the study (M1: 11kgs, 12 years; M2: 8,5kgs, 13 years). The project was authorized by the French Ministry for Higher Education and Research (# 2016120910476056 and #1588-2015090114042892) in accordance with the French transposition texts of Directive 2010/63/UE. This authorization was based on an ethical evaluation by the French Committee on the Ethics of Experiments in Animals (C2EA) CELYNE registered at the national level as C2EA number 42.

### Surgery

The animals were implanted with a peek MRI-compatible headset covered by dental acrylic. The anesthesia for the surgery was induced by Zoletil (Tiletamine-Zolazepam, Virbac, 5 mg/kg) and maintained by isoflurane (Belamont, 1–2%). Post-surgery analgesia was ensured thanks to Temgesic (buprenorphine, 0.3 mg/ml, 0.01 mg/kg). During recovery, proper analgesic and antibiotic coverage was provided. The surgical procedures conformed to European and National Institutes of Health Guidelines for the Care and Use of Laboratory Animals.

### Fluoxetine preparation

Fluoxetine hydrochloryde is a SSRI which binds to the human 5-HT transporter with a Ki of 0.9 nmol/l and is between 150- and 900-fold selective over 5-HT1A, 5-HT2A, 5-HT2c, H1, α1, α2-adrenergic, and muscarinic receptors (Ambati et al., 2021). The fluoxetine (N-Methyl-3-[(4-trifluoromethyl) phenoxy]-3-phenylpropylamine hydrochloride) used in the present study has a molecular weight of 345,78 g/mol. Powder galenic form (BioTechne©, ToCris BioScience) was diluted in a saline vehicle (NaCl) as follows. In order to inject the smallest possible volume to the monkeys, we dissolved fluoxetine in a saline solution at a concentration of 8 mg/mL, vortexed 10 seconds and heated the suspension at 60°C in bain-marie to increase solubility while not degrading the active compound. When needed, this preparation was frozen at -20°C so as to avoid the molecule degradation and heated back to body temperature when necessary.

### Fluoxetine administration

In order to reduce the stress potentially induced by the injection, monkeys were progressively trained to spontaneously receive subcutaneous saline injections with clicker training. In contrast with intramuscular injections, subcutaneous injections allow a slow distribution of injected product, thus a longer half-life in the body. Injection site and side of injection was changed daily. Injection sites were carefully monitored and sanitized. Once animals reached stable performance and were habituated to subcutaneous injections, behavioral data collection started under either placebo (saline) or fluoxetine injections (2,5mg/kg/day). This Fluoxetine concentration was chosen based on its specific serum concentration decay time in macaques (half-life <16h, Sawyer and Howell, 2011) and the reported threshold for behavioral effects (Fontenot et al., 2009; Chen et al., 2012). Two different injection schedules were used. Acute schedule involving, over a full working week, one day of saline injection, followed by one day of fluoxetine, followed by three days of saline injections (Free choice task). Chronical injections involved daily fluoxetine injections during one full month (Luminance perceptual task and target detection task under spatial uncertainty). The effect of this latter schedule was compared to that of one month of saline injections. Monkeys were always injected in the morning, at the same time, and behavioral data was collected 4 to 6 hours later based on the pharmacokinetics of fluoxetine (Sawyer and Howell, 2011). Table 1 describes the number of placebo and fluoxetine sessions collected for each task, under each injection schedule, as well as general trial statistics.

**Table 1.** Description of number of sessions and trial statistics. Number of sessions and trial statistics (median + se) are described per type of tasks (luminance detection task, target detection task with distractors and saccadic reward competition task), condition (Placebo and Fluoxetine) and monkeys. Injections for the saccadic reward competition task, were performed acutely (one day per week). For the other tasks, injections were performed according to a chronic injection schedule.

|  | **Monkey 1** |  |  |  | **Monkey 2** |  |  |  |
| --- | --- | --- | --- | --- | --- | --- | --- | --- |
|  | **Placebo** |  | **Fluoxetine** |  | **Placebo** |  | **Fluoxetine** |  |
|  | **Acute** | **Chronic** | **Acute** | **Chronic** | **Acute** | **Chronic** | **Acute** | **Chronic** |
| **Luminance detection task** |  | # Sess. =3;<br>Med. # tr.=<br>1257 +/-<br>233.98 |  | # Sess. =4;<br>Med. # tr.=<br>1914 +/-<br>13.96 |  | # Sess. =5;<br>Med. # tr.=<br>289 +/-<br>62.33 |  | # Sess. =5;<br>Med. # tr.=<br>387 +/-<br>14.81 |
| **Target**<br>**detection task**<br>**with**<br>**distractors** |  | # Sess. =8;<br>Med. # tr.=<br>970 +/-<br>72.85 |  | # Sess. =4;<br>Med. # tr.=<br>1443 +/-<br>69.01 |  | # Sess. =15;<br>Med. # tr.=<br>271 +/-<br>44.04 |  | # Sess. =10;<br>Med. # tr.=<br>497 +/-<br>86.39 |
| **Saccadic**<br>**reward**<br>**competition**<br>**task** | # Sess. =7;<br>Med. # tr.=<br>936 +/-<br>109.10 |  | # Sess. =7;<br>Med. # tr.=<br>1128 +/-<br>94.99 |  | # Sess. =7;<br>Med. # tr.=<br>624 +/-<br>87.64 |  | # Sess. =7;<br>Med. # tr.=<br>792 +/-<br>94.99 |  |

### Experimental setup

Monkeys sat in a primate chair in sphinx position head-fixated thanks to a surgically implanted head post. They were positioned in front of a screen. The eye to screen distance was of 60cm and screen resolution was 1200×1900pixels with a 60Hz refresh rate. Gaze location was sampled at 120Hz using an infrared video-eye tracking system (ISCAN). Eye Movement data Acquisition Software interfaced with an inhouse program for stimulus delivery and experimental control (Presentation©). Monkey hand responses were produced by releasing a bar, the effect of which was to restore the continuity of an infra-red optic beam.

### Behavioral tasks

Animals had free access to food and were maintained under a water regulation schedule individually optimized to keep a stable motivation and performance. They were trained on three different behavioral tasks. In all of these tasks, monkeys had to fixate a central fixation point on a screen for a variable duration (1-to-2 secs) while stimuli (size: 0.5°; duration: 100ms) were presented at an eccentricity of 8°.

#### Luminance detection task

This task aims at assessing changes in luminance perception thresholds under fluoxetine as compared to saline placebo injections. Monkeys had to fixate a central cross. One thousand to 2000ms from fixation onset, a 200ms target appeared randomly at one of the four possible following positions: (6√2, 6√2), (−6√2, 6√2), (−6√2, -6√2) or (6√2, -6√2). This task was designed to be dominated by bottom-up perceptual processes as we did not use any spatial cue to indicate to the monkeys the position of the upcoming target (Reynolds et al., 2000; Carrasco et al., 2004; Ibos et al., 2009; Reynolds and Chelazzi, 2004). The luminance of the target varied from the background luminance, from easy to hard, on a scale of seven equidistant luminance values (figure 1). Each target position was sampled for each luminance 10 times per session. Monkeys were rewarded for producing a hand response to target presentation, within a response time window of [150ms-1000ms]. Misses or false alarms are not rewarded. Note that by construction the task does not produce correct rejections. This task was tested twice, in two sets of recording sessions spaced by ten months and a wash out period of at least two consecutive months in between (first data collection: M1: 7 sessions, M2: 10 sessions; second data collection: M1: 3 sessions, M2: 4 sessions (table 1). Individual psychometric luminance perception curves are constructed for each of the four target positions independently for the placebo and the fluoxetine conditions. In the results, we discuss the data collected during the first data collection sessions. The data from the second data collection sessions are presented in supplementary material (figure S1). They are not significantly different from those reported for the first sessions, indicating a stable and reproducible effect in time.

#### Target detection task in the presence of distractors

The previous task allows to identify possible changes in individual perception thresholds. However, changes in such metrics can be due to bottom-up changes in perceptual sensitivity (or dprime) or to changes in individual subject response criterion. In order to refine our understanding of the effect of fluoxetine on perception and decision-making, we used a peripheral target detection task in the presence of spatial distractors (Figure 2). Monkeys had to fixate a central cross. One thousand to 2000ms from fixation onset, a 200ms target appeared randomly at one of the four possible following positions: (6√2, 6√2), (−6√2, 6√2), (−6√2, - 6√2) or (6√2, -6√2). Target luminance was defined as target luminance associated with a 70% correct detection threshold in the placebo sessions of the luminance task. Prior to actual target presentation, a 200ms spatial distractor could randomly appear within of virtual circle of 2° around the expected target location (as learned from the previous task). Distractors were identical to the target and only differed in their position. Distractors were present in 3:4 of the trials. Monkeys were rewarded for producing a hand response to target presentation, within a response time window of [150ms-1000ms]. Data on this task were collected from 8 (M1) & 15 (M2) placebo sessions and 4 (M1) & 10 (M2) fluoxetine sessions (table 1).

#### Saccadic reward competition task

In order to investigate the possible contribution of fluoxetine to the implementation of reward biases and learning, we used a saccadic competitive task, towards stimuli the spatial position of which was associated, with a specific reward probability schedule (Figure 3). The specific spatial reward contingencies changed from one session to the next. Monkeys had to fixate a central cross. One thousand to 2000ms from fixation onset, two identical stimuli were presented. Stimuli were drawn from a virtual array of eight stimuli organized along a circle of 8° of eccentricity. From one session to the other, each location in this virtual array was associated with a different reward probability (stable across trials of the same session), which the monkeys discovered at the beginning of the session, thus building a reward based spatial priority map (Chelazzi et al., 2014; Della Libera et al., 2017), then exploited during the rest of the session. Possible high reward probabilities were 80%, 50% and 20%, according to a fixed spatial relationship, such that the extreme reward probabilities (80% and 20%) were neighbored intermediate reward probability targets (50%) (figure 3). Monkeys had to make a saccade to one of the two presented stimuli and were rewarded according to the reward probability associated with the chosen target location. The spatial reward contingency map was rotated from one day to the next, leading to seven different spatial reward contingency maps, played several times over independent sessions (as the initial spatial contingency map on which initial training was performed was not used). For this experiment, we used a 3-week chronic saline injection schedule followed by and 8-week chronic fluoxetine injection schedule.

### Data analysis

All analyses are implemented in Matlab® using ad-hoc scripts.

#### Extracted behavioral and physiological measures

For each task and each session, we quantified *<u>session length</u>* (overall number of trials) and overall *<u>behavioral performance</u>* (percentage of correct trials relative to the sum of correct and miss trials). For the luminance detection task, we computed the behavioral performance independently for each target contrast level to right and left targets. We then fit a sigmoid model to the data. Using a sigmoid function (R P (2022). Sigm_fit (https://www.mathworks.com/matlabcentral/fileexchange/42641-sigm_fit), MATLAB Central File Exchange. Retrieved August 30, 2022), we determined the *<u>p50</u>* (target luminance associated with a 50% chance performance), the *<u>slope</u>* (sensitivity to contrast changes) and the *<u>baseline</u>* (response level to noise) for each mean number of trials per session (see table 1), in both placebo and fluoxetine condition. On the two target detection tasks, *<u>manual reaction times</u>* (RT) were extracted, defined as the time between target presentation and hand lever release (Figure 6, for RT distributions). Pupil size variation (McGuirk and Silverstone, 1990; Phillips et al., 2000; Dumont et al., 2005) and blinks were identified (Wilson et al., 1983; Semlitsch et al., 1993) and we quantified, for each session, the distribution of *<u>blink duration</u>* and *<u>pupil size variations</u>* (Figure 5). We quantified durations of all blinks while monkeys where engaged in tasks in both placebo and fluoxetine conditions and determined the median value. As for measuring pupil dilation, we estimated pupil size during a period of 3 seconds of rest before task initiation while monkeys were sat in the dark, and during a period of 3 seconds during task execution, sampled after target presentation. Percentage pupil size variation in placebo and fluoxetine conditions were computed by estimating the average pupil size on these two epochs and normalizing it by average pupil size over the entire task duration.

#### RT analyses

All RTs above 1000ms were excluded from the analysis. RT distributions were then analyzed using the Later model (Noorani and Carpenter, 2016). This model distributes data according to their frequency of distribution. The LATER model allows to segregate RTs in two categories: controlled and anticipated/express responses. We thus segregated, for each task, RT as a function of target position, and we used the LATER model analysis in order to identify the cutoff between anticipatory and controlled responses (Table 2).

**Table 2.**
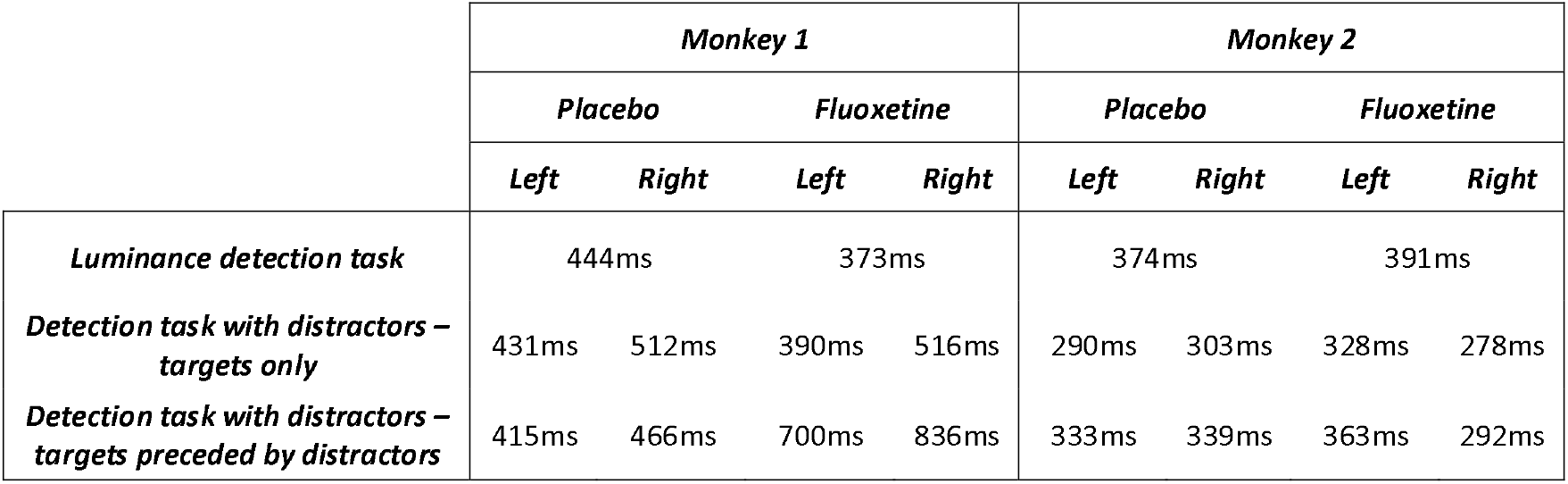
RT threshold between anticipatory and controlled responses as defined by the LATER model. This threshold was defined independently for each monkey, each task and each target position, except for the luminance detection task, for which all four positions were considered together.

#### Signal Detection Theory

In the target detection task in the presence of distractors, we used signal-detection-theory and computed the monkey’s sensitivity to the location of the target relative to the distractors randomly presented around the target location (d’, reflecting bottom-up sensory features) as well as their response criterion (reflecting top-down control in the decision-making process). These metrics were independently computed per session (see table 1).

#### Spatial (SSI) and reward (RSI) selectivity index

In the saccadic reward competition task, we calculated, for each session (Table 1), the choice performance of a given singleton for each possible pair of stimuli. We then calculated, for each session, and each spatial position the SSI as follows. For each position i, we computed the median choice percentage SSI_i_ that a singleton at position i was chosen, irrespective of the second singleton in the pair and irrespective of their associated rewards. In other words, it is the median, over all pairs containing singleton i, of the percentage of times I was actually chosen. SSIi thus reflects the average preference of the monkey for singleton i.

Likewise, we computed the RSI as follows. For each reward contingency i, we computed the median choice percentage RSI_i_ that a singleton with that specific reward contingency i was chosen, irrespective of the reward associated with the second singleton in the pair, and irrespective of their spatial position. RSI_i_ thus reflects the average preference of the monkey for a given reward contingency i.

### Statistical analyses

All statistical analyses are non-parametric Wilcoxon or Kruskall-Wallis tests, except when two-way ANOVAs are required. *P*-valuel⍰<⍰0.05 were considered as statistically significant. All statistical analyses are implemented in Matlab® using ad-hoc scripts.

## Data availability

The data that support the findings of this study are available from the corresponding author upon reasonable request. A Source Data file provides the raw data used to create all of the figures of this paper.

## Code availability

The code that supports the findings of this study is available from the corresponding author upon reasonable request.

## Acknowledgements

With financial support from UNADEV (National Union for Blind and Visually Impaired People) in partnership with ITMO NNP (Multi-Organism Thematic Institute Neurosciences, cognitive sciences, neurology and psychiatry) / AVIESAN (national alliance for life science and health) as part of research on vision disorders to S.B.H. as well as LABEX CORTEX funding (ANR-11-LABX-0042) from the Université de Lyon, within the program Investissements d’Avenir (ANR-11-IDEX-0007) operated by the French National Research Agency (ANR). We thank Fidji Francioly and Laurence Boes for animal care. We thank Serge Pinède and Julian Amengual for technical assistance on the project.

## Author Contribution Statement

Conceptualization, S.B.H., M.G; Stimuli preparation, M.G; Data Acquisition, M.G.; Methodology, M.G., S.B.H.; Investigation, M.G., S.B.H.; Writing – Original Draft, M.G., S.B.H.; Writing – Review & Editing, M.G., S.B.H.; Funding Acquisition, S.B.H.; Supervision, S.B.H.

## Competing interest statement

The authors declare no competing interests.

## Ethics declaration

Animal experiments were authorized by the French Ministry for Higher Education and Research (project no. 2016120910476056 and 1588-2015090114042892) in accordance with the French transposition texts of Directive 2010/63/UE. This authorization was based on ethical evaluation by the French Committee on the Ethics of Experiments in Animals (C2EA) CELYNE registered at the national level as C2EA number 42.

## Supplementary material

**Figure S1:**
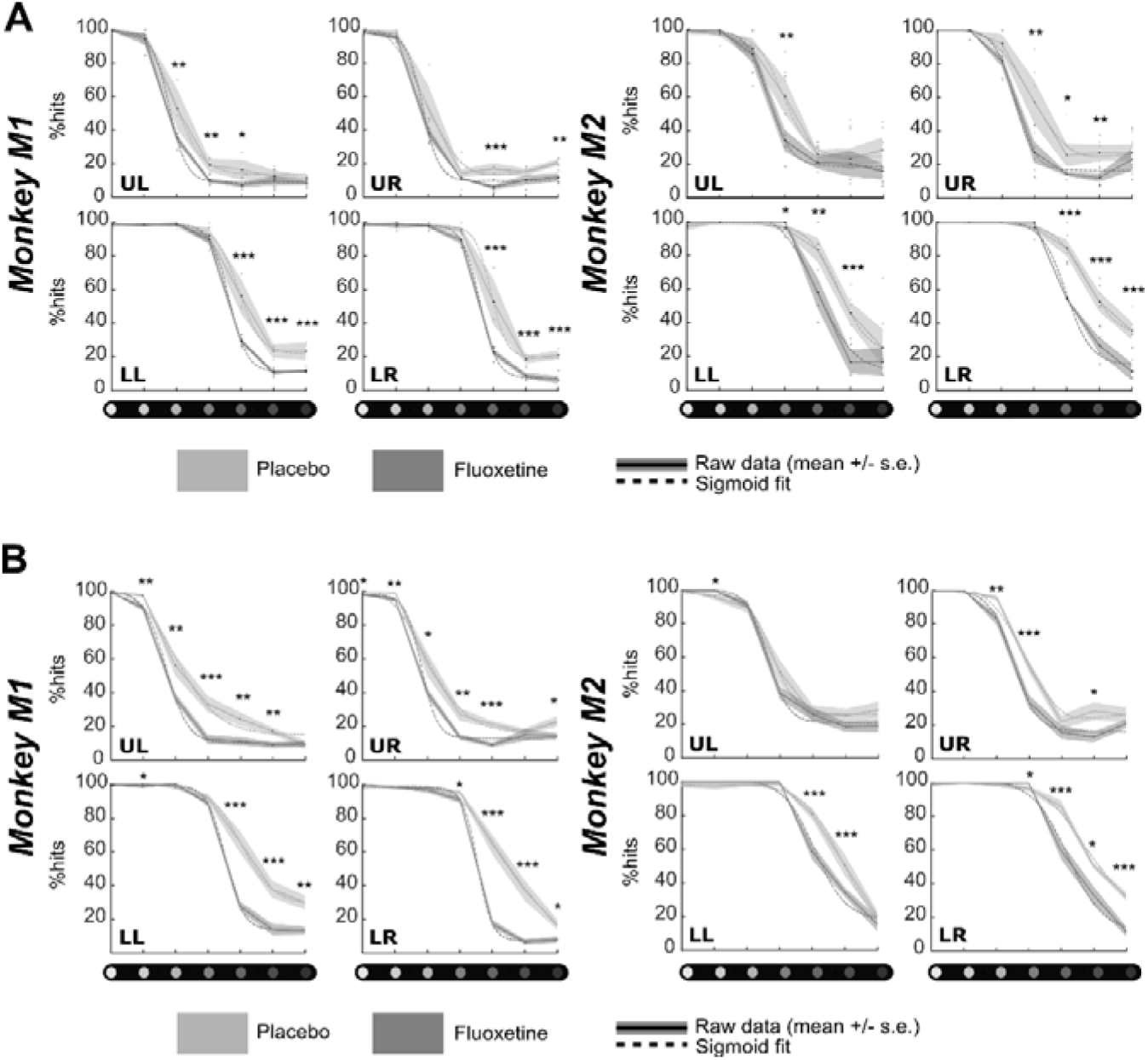
Effect of Fluoxetine on perceptual thresholds in a luminance detection task. For both monkeys, % of hits were computed independently for each target luminosity. Dots represent individual sessions, continuous lines represent average % hits across all sessions (+/-s.e.) and dashed lines represent sigmoid fit of the data. Placebo data are represented in light gray and Fluoxetine data are represented in dark gray. Behavioral data are represented independently for each target position. Statistical significance is represented as follows: ***, p<0.001; **, p<0.01; *, p<0.05; n.s., p>0.05. (A) First experiment (same data as in figure 2). (B) Second identical experiment at 10 months’ interval.

**Figure S2:**
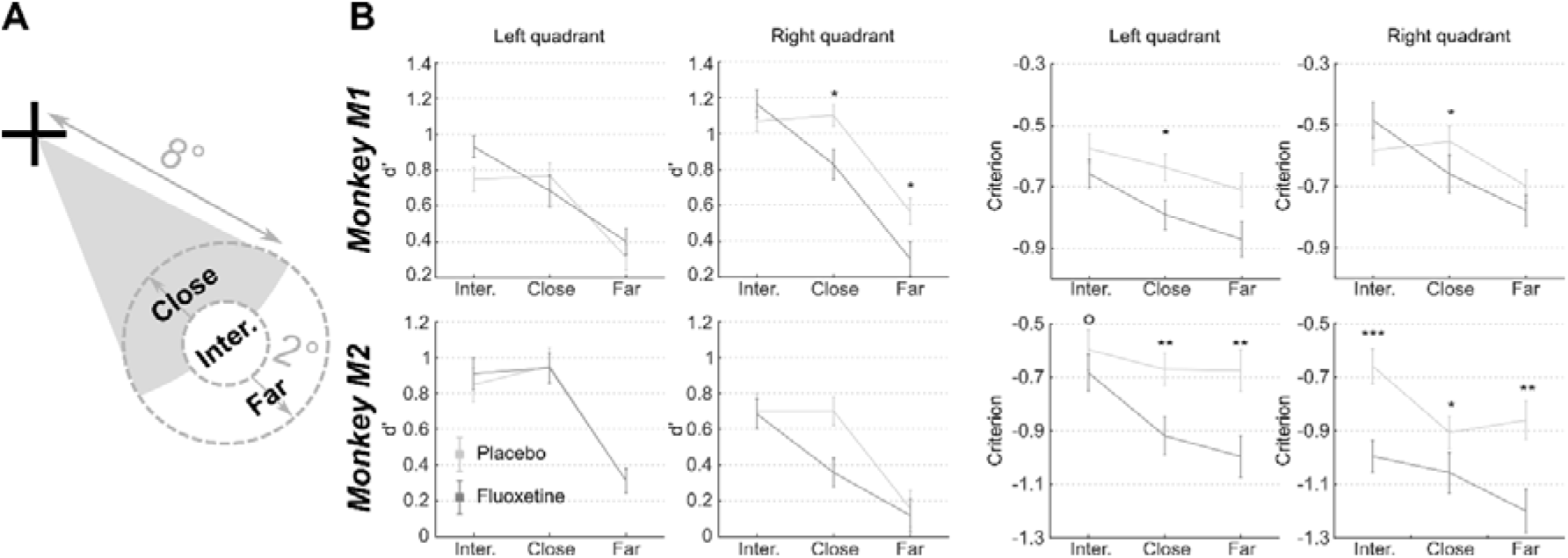
Effect of Fluoxetine on spatial sensitivity d-prime and response criterion in a target detection task in the presence of spatial distractors, as a function of target distractor distance. (A) Spatial categorization of target to distractor distances. (B) For both monkeys, d’ and criterion were computed independently for each target and for each target to distractor distance. Median +/-s.e. of median are presented for placebo and Fluoxetine conditions. Statistical significance is represented as follows: ***, p<0.001; **, p<0.01; *, p<0.05; °, p<0.07. Figure 3 represents this data irrespective of target to distractor distance.

**Figure S3:**
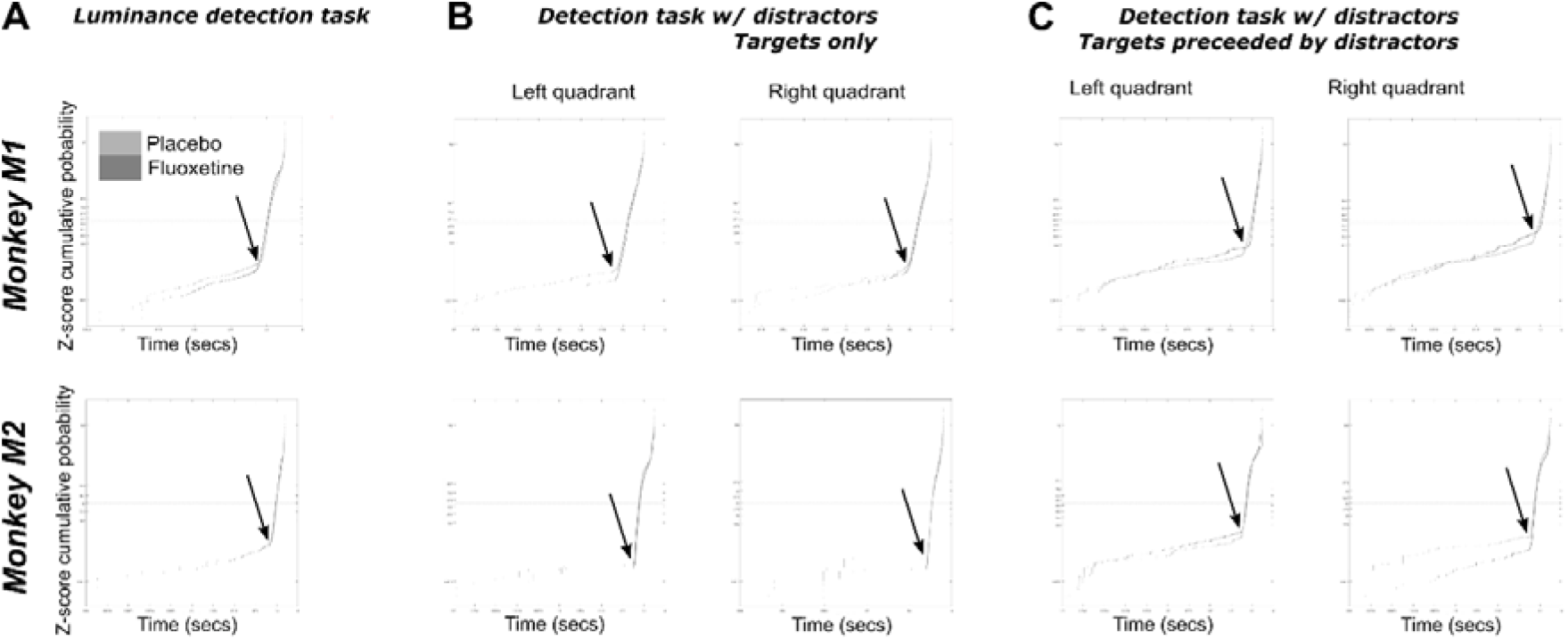
Identification of limit between anticipatory and controlled RT using the LATER model, for monkeys M1 (top) and M2 (bottom), in the placebo (light gray) and Fluoxetine (dark gray) conditions in the luminance task (A, all positions collapsed, high luminance targets only), in the detection tasks with distractors, on target only trials (B), and in trials with distractors (C).

## Notes

### Competing Interest Statement

The authors have declared no competing interest.

